# A loss of crAssphage stability in the human gut virome is associated with obesity and metabolic syndrome

**DOI:** 10.1101/2022.06.15.496347

**Authors:** Melany Cervantes-Echeverría, Luigui Gallardo-Becerra, Fernanda Cornejo-Granados, Adrian Ochoa Leyva

## Abstract

Viral metagenomics studies of the human gut microbiota unravel differences in phage populations between healthy and disease, stimulating interest in the role that phages play in bacterial ecosystem regulation. CrAssphages are not only the most abundant viruses but also are a common component of the gut phageome across human populations. However, the role of crAssphages in obesity (O) and obesity with metabolic syndrome (OMS) remains largely unknown. Therefore, we explored the role that crAssphages have on both diseases in a children’s cohort. We found decreased crAssphage abundance, prevalence, richness, and diversity in O and OMS compared to normal-weight (NW), suggesting a loss of crAssphages stability in the human phageome associated with the disease. Interestingly, when we analyzed the abundance of host crAssphages bacteria, we found that Bacteroidetes, Bacteroidia, and Bacteroidales were significantly decreased in O and OMS, suggesting a possible relation with the loss of crAssphages stability. Regarding crAssphage taxonomy, a significantly decreased abundance of the crAssphage Alpha subfamily and the Alpha_1 and Alpha_4 genus and a significant overabundance of the Delta_8 was found in OMS. A strong taxonomical signature of obesity is the over-abundance of Bacilli, which also were significantly increased in O and OMS. Notably, we found a significant negative correlation between crAssphages and Bacilli abundances, suggesting an association between the decreased abundance of crAssphage and the over-abundance of Bacilli in OMS. Furthermore, we found a loss of crAssphage stability in the human virome associated with the presence of obesity, having a more significant impact on obesity with metabolic syndrome, suggesting that these bacteriophages could play an essential role in inhibiting metabolic syndrome in obese individuals. Our results open a promising treatment for these diseases through fecal crAssphage transplantation.

## Introduction

Recent advances in bioinformatics and experimental protocols for viral metagenomics showed us that the gut virome is mainly dominated by bacteriophages (Bikel et al., 2022, Shkoporov & Hill, 2019). Indeed, there are disease-specific changes in the gut virome in inflammatory bowel disease (Norman et al., 2015; Zuo et al., 2019), AIDS (Monaco et al., 2016), diabetes (Ma et al., 2018), malnutrition (Reyes et al., 2015), obesity, and obesity with metabolic syndrome (Bikel et al., 2021). There are many reasons to study the relationship between phages and microbiota, including that they kill specific bacteria and transfer different genes; consequently, they can alter the host’s relationship with the microbiota (Rossi et al., 2020). Human feces have been estimated to contain 10e9 VLPs per gram of feces (Bikel et al., 2021). Although also there are “passenger viruses” that are diet-associated, such as plant and dietary viruses (Minot et al., 2011).

Since their discovery in 2014, crAssphages are one of the most intriguing elements of the human virome (Dutilh et al. 2014, Yutin et al., 2017), being one of the most abundant gut associated-virus and also is a common component of the gut virome across the human population (Edwards et al. 2019, Cervantes-Echeverría et al., 2018), showing a considerable genomic diversity (Guerin et al., 2018). Furthermore, the crAssphages appear to have coevolved with humans (Edwards et al. 2019). Indeed, crAssphages are associated with several non-human primates populations globally distributed (Edwards et al. 2019). The crAssphages also appear to be abundant and widespread in diverse animal and environmental-human habitats (Yutin et al., 2018). Several bacteria of the phylum Bacteroidetes appear to be the primary hosts of crAssphages, being the 0CrAss001, the first crAssphage member isolated and infecting the Bacteroides intestinalis (Shkoporov et al., 2018). Recently, the taxonomy for crAssphages composed of four subfamilies and ten genera was proposed (Guerin et al., 2018). Indeed, despite the high interindividual variations in the gut virome (Bikel et al., 2021), the crAssphages constitute a global virus marker for global-scale studies (Edwards et al., 2019), and it is part of the human virome core (Stockdale et al., 2021).

The crAssphages are generally scarce in the feces of one-month infants, increasing their abundance in the four-month and the 2-5-year-old infant and becoming a high-level persistence of crAssphages in adults (Gregory et al., 2020). Indeed, these phages are detected in both the mother and the infant, suggesting vertical transmission, and they also can be acquired through fecal microbiota transplantation (Siranosian et al., 2020). These discoveries have stimulated the study of crAssphage phages as an indicator of human fecal pollution (Cinek et al., 2018; Stachler et al., 2017).

Recent research suggests that crAssphage prevalence is associated with an industrialized lifestyle/diet but with no associations with health, age, sex, or body-size variables (Honap et al., 2020, Edwards et al., 2019). CrAssphage abundance was also not associated with diseases such as diarrhea, CD, HIV, IBD, T2D, and malnutrition (Liang et al., 2016, Guerin et al., 2018, Honap et al., 2020). However, a recent study showed that the crAssphage was more abundant in the healthy controls than in colorectal cancer, suggesting a potential treatment strategy for this disease through fecal CrAssphage transplantation (Gao et al., 2021). On the contrary, the abundance of crAss-phage was significantly higher in UC patients than in the normal control (Yang et al., 2020). Nonetheless, as significant components of the human gut, crAssphages deserve more attention and the association with specific diseases shall be further investigated. No studies address the crAssphages role in obesity and metabolic syndrome. The study of this expansive group of viruses is still in its infancy, and fundamental features of these major players in the human virome and their actual spread in the biosphere remain to be studied (Stockdale et al., 2021).

In this study, we explored the role that crAssphages have in obesity and metabolic syndrome to understand their biological significance better as a critical component of the human gut virome. Further, we evaluated crAssphage abundance as a function of gut bacteria, anthropometric and biochemical parameters typically altered by obesity and metabolic syndrome.

## Results

### 1. Different abundance and prevalence of crAssphages were detected at the same sequencing depth

We used our viral metagenomics sequencing data of 28 fecal samples from a previously described cohort of 7- to 10-year-old children, ten with normal-weight (NW), ten with obesity (O), and eight with obesity and metabolic syndrome (OMS) (Bikel et al., 2021) (Table S1).

To use the same sequencing depth for all samples, we generated 1,000 sets of random sub-samplings of 700,000 paired reads per sample (see methods). First, only seven samples from the NW group, nine from group O, and six from the OMS group met the sequencing depth requirements to be subsampled (Table S2). Second, each set of subsampled reads was mapped against the crAssphages genome database composed of 280 non-redundant genomes (Table S3, see methods). In this manner, we obtained the number of mapped reads for each crAssphage genome as the average of the 1,000 sets per sample. Finally, this average normalized read-counts table was used for further analysis.

From the 280 crAssphages, only 208 genomes contained mapped reads, and their relative abundance ranged from 0.0011 to 3.654% of total reads per sample (Table S4). We found a tendency to decrease the crAssphage abundance in obesity with metabolic syndrome compared to NW and O (Figure 1a). However, the difference was not significant. In addition to decreased abundance, the prevalence was also decreased in O and OMS (Figure 1b). Again, however, the difference was not significant.

**Figure 1.**
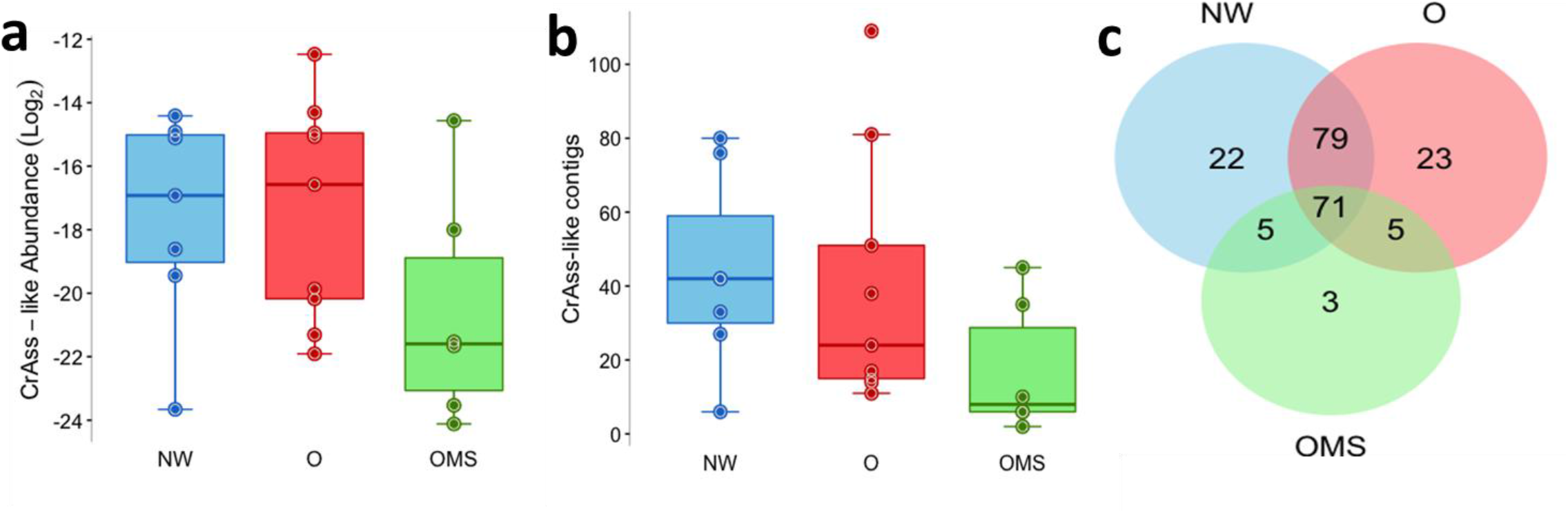
crAssphage abundance and prevalence. a) abundance of crAssphage genomes. Points represent the average abundance for each sample, with boxes showing the group’s distribution. b) prevalence of crAssphage genomes. Points represent the average prevalence for each sample, with boxes showing the group’s distribution. c) Venn diagram of the presence of crAssphage genomes between groups. Error bars represent the mean ± SD.

We observed that 71 phages were shared among the three groups, and several phages were unique for each group (Figure 1c). The distributions of crAssphages present in each sampl are shown in Figure 2.

**Figure 2.**
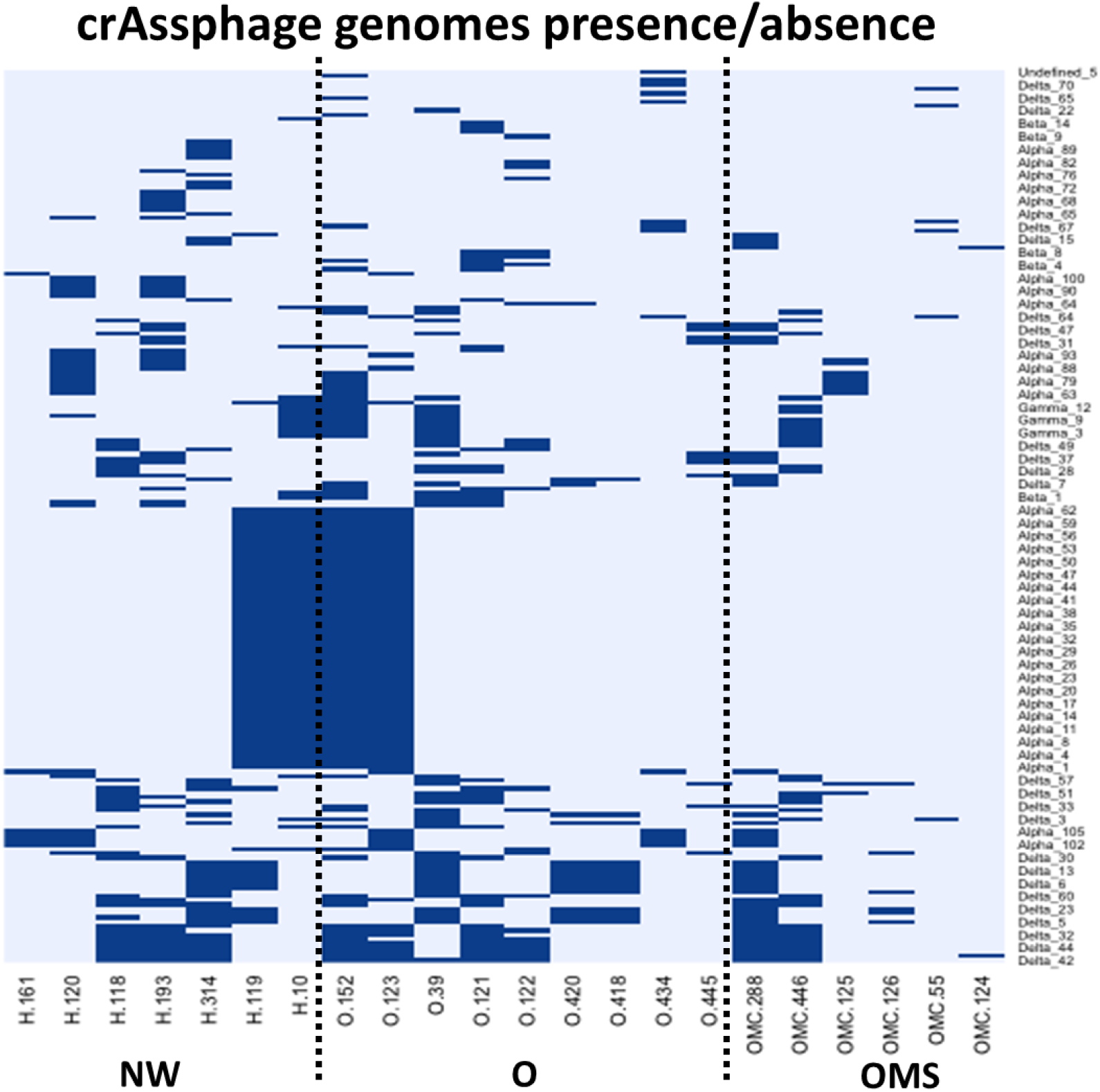
Presence-absence heat map of crAsphage genomes distribution among all samples.

### 2. Changes in crAssphage taxonomy characterize diseases

Using the normalized reads-count table, we calculated the relative abundance of the four subfamilies and ten genera described by Guerin et al. 2018. The Alpha and Delta subfamilies were the most abundant among the three groups (Figures 3a and 3c). The Alpha was significantly (p = <0.01) enriched in NW (mean= 0.00592) as compared to O (mean = 0.00274) and OMS (mean = 0.00148). The Beta family was significantly (p = <0.01) enriched in OMS (mean = 0.00298) as compared to O (media= 0.00040), however when was compared to NW (mean = 0.00006) the difference was not significant. An over-abundance of Delta was also observed in OMS (mean = 0.01126) and O (mean = 0.00989) as compared to NW (mean = 0.00529); however the difference was not significant.

**Figure 3.**
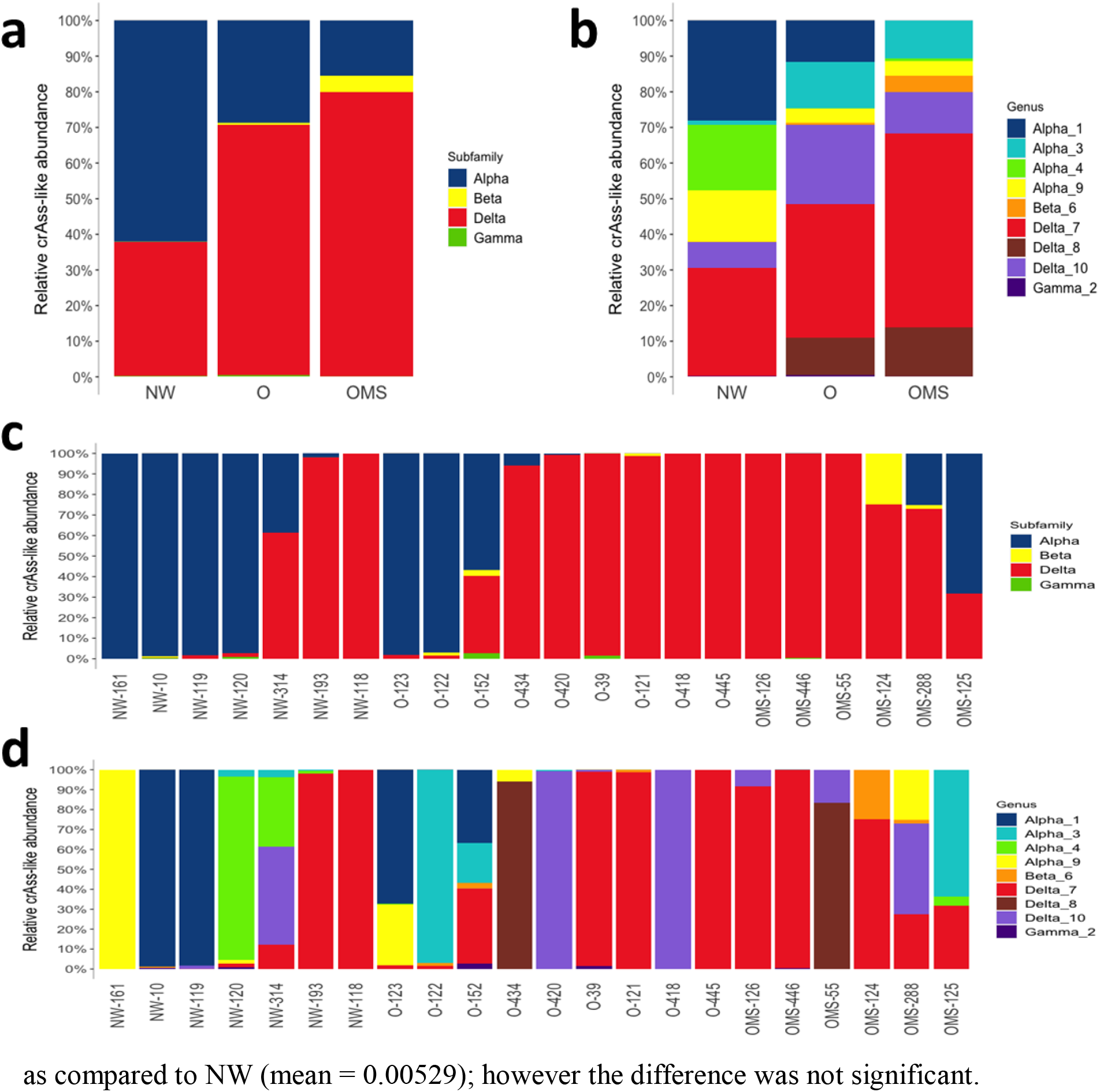
Relative abundance of crAssphage genomes grouped by taxonomy. a) the average of subfamilies and b) the genus per group, c) the average of subfamilies, and d) the genus per sample.

At the genus level the Delta-7 represented the most abundant phages among the three groups; however no significant difference between groups was observed (Figures 3b and 3d). Contrary, the Alpha_1 was significantly (p a= >0.01) increased in NW (mean = 0.00454) as compared to O (mean = 0.00187) and OMS (mean = 0), and Alpha_4 also was significantly (p = >0.01) increased in NW (mean = 0.01013) as compared to O (mean = 0.00003) and OMS (mean = 0.00047). The Beta_6 was significantly over-abundant in OMS (mean = 0.00330) as compared to O (mean = 0.00040), however when was compared to NW (mean = 0.00006) the difference was not significant. The Delta_8 was significantly (p = >0.01) over-abundant in OMS (mean = 0.01263) as compared to O (mean = 0.00953) and NW (mean = 0). Interestingly, phages from genus Gamma_5 were not present in any sample, while Delta_8 and Alpha_1 were depleted in NW and OMS, respectively.

### 3. Decreased crAssphages diversity and richness are associated with the disease

The normalized reads-count table was used to analyze the alpha diversity among samples (see methods). The alpha diversities showed that phage diversity and richness decreased in O and OMS compared to NW (Figures 4a and 4b), although the differences among groups were not significant. We also performed a beta-diversity analysis to know if crAssphages clustering was associated with the disease; however, no clustering was observed (data not shown).

**Figure 4.**
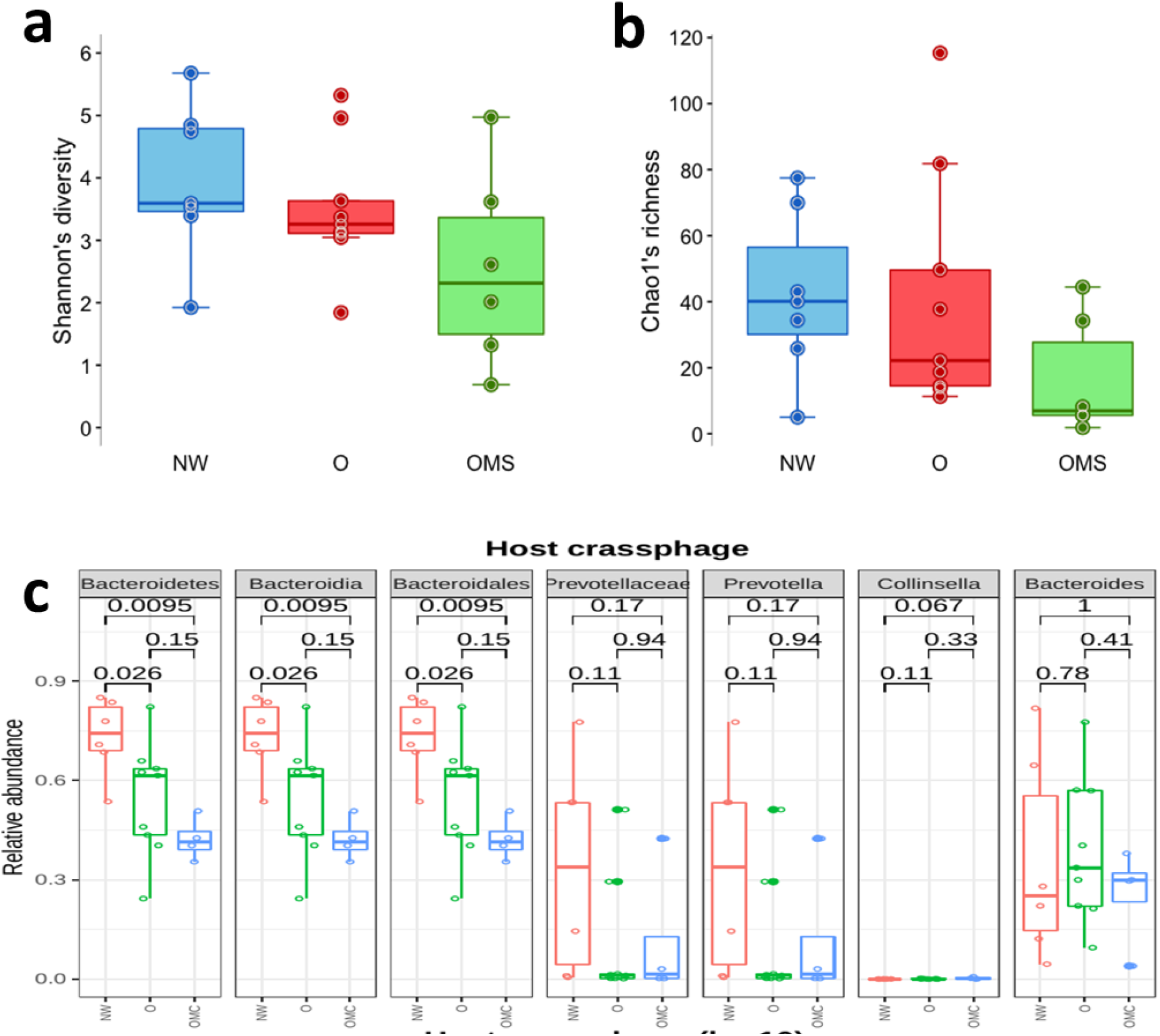
Alpha diversity of crAssphage genomes and relative abundance of bacterial crAssphage hosts. a) crAssphage diversity. Each point shows the mean of normalized read-counts table Shannon diversity for each sample, with boxes showing the group’s distribution group. b) crAssphage richness. Each point shows the mean of normalized read-counts table observed genomes for each sample, with boxes showing the group’s distribution group. Differences in (a) and (b) were not significant. Error bars represent the median ± SD. c) relative abundance of bacterial crAssphage hosts. Each point shows the abundance for each sample, with boxes showing the group’s distribution group. Error bars represent the median ± SD. The p-value resulting from comparisons among groups is shown above the box plots.

### 6. Host bacteria of crAssphages were found to decrease in the disease

We analyzed the abundances of the seven taxa suggested as crAssphage hosts among the three groups (Figure 4c). We found that Bacteroidetes, Bacteroidia, and Bacteroidales were significantly decreased in O and OMS compared to NW (Figure 4c). We next assessed whether the abundance of the total crAssphages was associated with a parallel change in suggested host bacteria. To this end, we selected the seven host bacteria taxa and calculated the Spearman correlation between their abundance and the crAssphages abundance in all samples. However, we do not find any significant correlation (data not shown). Although the correlations were not significant, could be an association between the significant decrease in Bacteroidetes, Bacteroidia, and Bacteroidales with the decreased abundance, richness, and diversity of crAssphages in O and OMS.

### 7. Increased Bacilli was associated with decreased crAssphage abundance in OMS

We also assessed whether all crAssphages were associated with parallel changes in bacterial populations previously associated with obesity and obesity with metabolic syndrome (Gallardo-Becerra et al., 2020). To this end, we calculated the Spearman correlation between the 27 bacterial taxa and crAssphages abundances in all the samples. Interestingly, we only found one significant correlation, a negative correlation between the crAssphages abundance and the order Bacilli (Figure 5a). Notably, Bacilli were also significantly increased in OMS compared to NW (Figure 5b).

**Figure 5.**
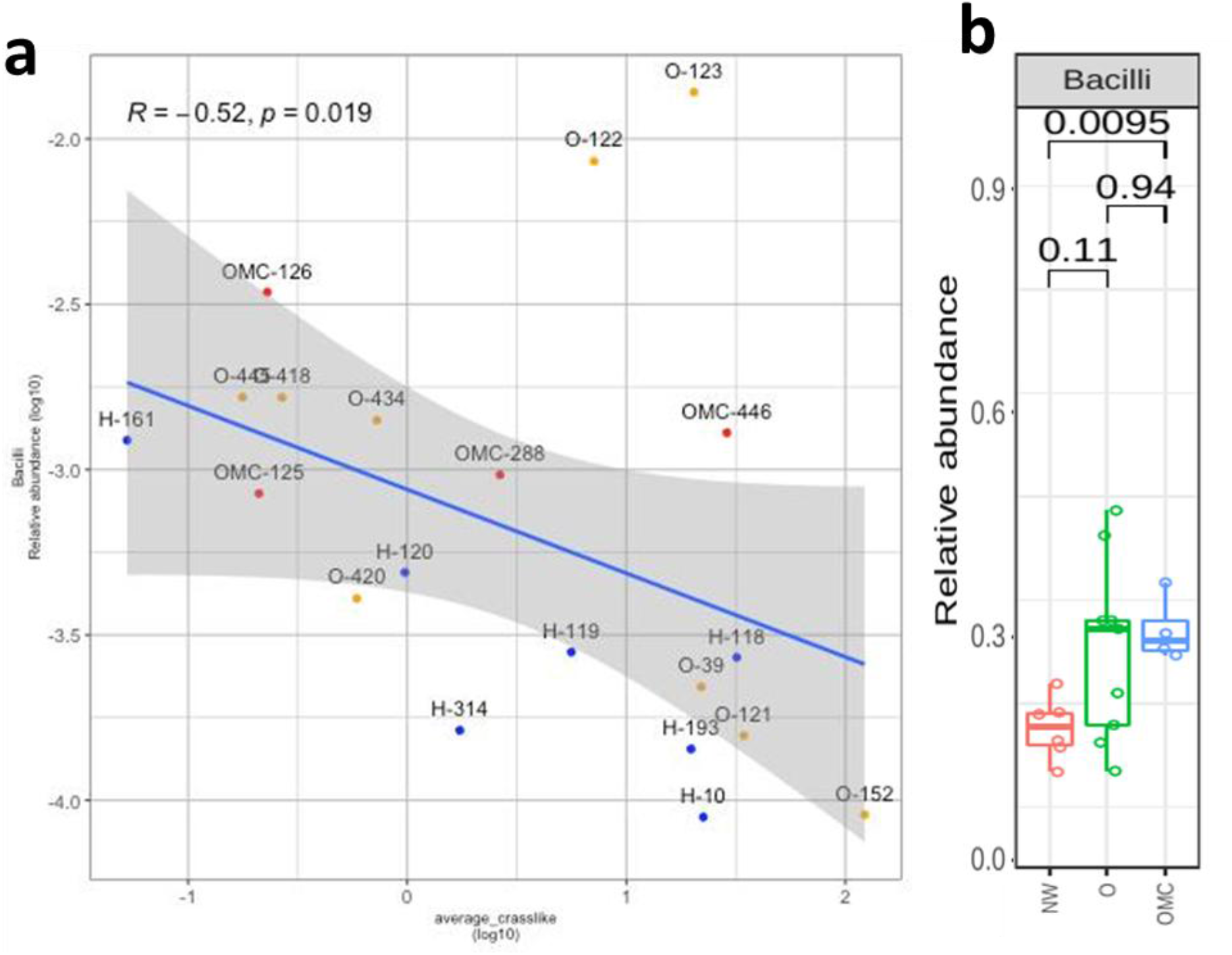
Relative abundance and Spearman correlation between Bacilli and crAssphage genomes. a) Spearman correlation plot between Bacilli and crAssphage abundances. Blue circles = NW samples; orange circles = O samples; and red circles = OMS samples. b) Relative abundance of Bacilli. Each point shows the abundance for each sample, with boxes showing the group’s distribution group.

We also analyzed whether crAssphage abundance correlated with the anthropometric and clinical parameters typically altered in obesity and metabolic syndrome, such as high body mass index (BMI), low levels of high-density lipoprotein (HDL), high levels of triglycerides, high glucose level, high waist circumference, and high weight (Gallardo-Becerra et al., 2020). However, we do not find any significant correlation.

## Discussion

The crAssphages have a central role in the gut virome because they are present in almost all human samples. The overall objective of this study was to gain insights into the potential role of crAssphages in children’s obesity and obesity with metabolic syndrome. The quantitative analysis of the crAssphage genomes revealed that most of our samples contained varying amounts of crAssphage genomes, from 0.0011 to 3.654% of total reads. Similar crAssphage abundances also were reported in healthy Malawian infants (Reyes et al., 2015).

Bacteroidetes have been proposed as the host phylum for crAssphages (Dutilh et al., 2014). Further investigations have also linked the Bacteroidales, Collinsella, Bacteroides, Bacteroidia, and Prevotellaceae as host bacteria. Specifically, two crAssphages have been isolated in pure culture, the ΦcrAss001 and ΦcrAss002, infecting Bacteroides intestinalis and Bacteroides Xylanisolvens, respectively (Shkoporov et al., 2018, Guerin et al., 2021). In this regard, we tested if our samples’ abundance of proposed host bacteria for crAssphages correlated with the crAssphage abundance. However, we do not find significant correlations between the abundance of crass and its hosts. This could be associated that crAssphages also replicate in a way that does not disrupt the proliferation of the host bacterium, maintaining itself in the continuous host culture for several weeks, suggesting their ability to maintain the stable colonization of the mammalian gut (Shkoporov et al., 2019).

Indeed, persistent propagation of virulent phages can occur in the human gut without host elimination (Guerin et al., 2021). Since there was no significant correlation between the abundance of bacterial hosts and the crAssphages, we cannot suggest that the loss of these phages’ abundance, diversity, and richness is a consequence of the loss of their hosts. However, it is essential to note that the data used for bacterial abundances were taken from 16S rRNA sequencing. Although a correlation between the abundance of crAssphage and Bacteroides dorei using 16S rRNA data was reported (Cinek et al 2017). Shot-gut sequencing could clarify these questions in future studies. Nonetheless, it is interesting that a significant decrease in the abundance of Bacteroidetes, Bacteroidia, and Bacteroidales was observed in obesity and metabolic syndrome, which could be associated with the decreased abundance, diversity, and richness also observed in crAssphages in obesity and metabolic syndrome.

A recent analysis found no significant correlation between crAssphage abundance in health and diseases in 1135 individuals. The study examined the correlation between the crAssphage abundance and 207 human variables, including 39 diseases (Edwards et al., 2019). Other studies showed no significant differences in the crAssphage abundance in fecal samples between diarrhea and healthy adults (Liang et al., 2016). In addition, we did not find any significant correlation between crass phages and the anthropometric and clinical parameters typically altered in obesity and metabolic syndrome.

Compared to normal control, a significant increase in crAssphage abundance was reported in patients with ulcerative colitis (Yang et al., 2020). Contrary, a recent study showed a decreased crAssphage abundance as a consequence of colorectal cancer compared to healthy controls, suggesting a promising treatment through fecal CrAssphage transplantation for this disease (Gao et al., 2021). In line with this, we found a decrease in abundance, richness, and diversity due to obesity and metabolic syndrome. Additionally, we also observed a decreased prevalence of crAssphages in the disease. A previous study of the human phageome also found a decreased abundance of crAssphages in obesity and metabolic syndrome compared to normal weight (Bikel et al., 2021). Our results also suggest a depletion of crAssphages in children’s viromes due to obesity and metabolic syndrome.

We found a loss of crAssphage stability in the human virome associated with the presence of obesity, having a more significant impact on the obesity with metabolic syndrome, suggesting that these bacteriophages could play an essential role in inhibiting the metabolic syndrome in obese individuals. Contrary, an increased richness and diversity for all the phageome was observed as a consequence of obesity and metabolic syndrome in the same children cohort. Further studies will be required to understand why the abundance, prevalence, diversity, and richness of crAssphages was decreased in obesity and metabolic syndrome, contrary to the observed for all phageome, as well as their significance for the human intestinal physiology and disease. The exact mechanism behind this trade-off needs further elucidation. A significantly increased abundance of Bacilli and its families Streptococcaceae and Lactobacillaceae was reported in obesity across two independent and large American adult populations (Peters et al., 2018). Notably, we found that Bacilli also was significantly over-abundant in OMS than in NW. Furthermore, a negative correlation was found between the abundance of crAssphages and the order Bacilli, suggesting that the decrease of crAssphages could be associated with the over-abundance of Bacilli.

The present study showed that the crAssphages were more abundant in the healthy control than diseased individuals, indicating a promising potential treatment strategy for obesity and metabolic syndrome through fecal crAssphage transplantation. In this regard, crAssphage is frequently transmitted via fecal microbiota transplantation (FMT) and can engraft stably in FMT recipients for up to one year (Siranosian et al., 2020). Furthermore, the crAssphages also could stably engraft and gradually dominate the viromes without affecting the levels of their host populations (Draper et al., 2018), offering an opportunity for treatment against the bacterial changes induced by obesity and metabolic syndrome (Gallardo-Becerra et al., 2020). Indeed, supplemental phage intake had no significant impact on overall health status and gut microbiota, and only specific bacteria were altered (Febvre et al., 2019). Also, a recent study suggests that phages may be applied as a dietary supplement in healthy and gastrointestinal distress individuals without causing exacerbation of symptoms (Gindin et al., 2018).

Additionally, crAssphage use an unusual strategy to establish themselves in the gut and to then persist stably within the microbial communities for several weeks and months (Shkoporov et al., 2019). Interestingly, changes in the abundance of crAssphages were reported after using traditional Chinese medicine (Yang et al., 2020), suggesting that crAssphage could also be modulated using pharmaceutical treatments. Therefore, further studies on the role of crAssphage in obesity and metabolic syndrome individuals should be investigated.

Our work also provides a significant differential abundance of specific crAssphage taxa associated with obesity for the first time. Specifically, the abundance of Alpha family and Alpha_1 and Alpha_4 genus were decreased in OMS, contrary to an overabundance of the Delta_8. Interestingly, phages from genus Delta_8 and Alpha_1 were depleted in NW and OMS, respectively. However, more studies will be required to determine the biological significance of these observations and whether this differential presence of specific crAssphage taxa was associated with obesity and metabolic syndrome development.

## Methods

### Viral Data acquisition and crAssphage genome mapping

The quality-filtered reads from gut metagenomic data of VLPs were obtained from a previous report by our group (Bikel et al. 2021). From the data set, 1,000 random sub-samplings of 700,000 paired reads were carried out per sample and used for further analysis to maintain the same sequence depth for all samples. Six samples were eliminated because they did not met this minimal number of reads. Thus, each set of reads from each of the 1,000 subsampling exercises per sample was mapped using SMALT to the reference crAssphage genome database consisting of 280 non-redundant crAssphage genomes, including 248 contigs previously validated by Guerin et al., 2018, 30 crAss-like contigs identified by Yutin et al., 2018, the Mexican-crAssphage genome (Cervantes-Echeverría et al., 2018) and the first crAssphage genome assembled by Dutilh et al., 2014 (Table S3). Additionally, we added five non-intestinal phage genomes as negative controls for the read mapping (Table S5). The parameters used were: 70 % of identity and a minimum of 60 nucleotides of the query read length covered by k-mer word seeds. All genomes were concatenated in one multifasta archive for the read mapping simultaneously. Finally, 22 samples were selected to continue with further analysis (Tables S2). In this manner, we obtained 1,000 tables of read mapping per sample, which were averaged to obtain a final table of read counts per sample. This table was then imported into R for abundance statistical analyses of crAssphages. Thus, the relative abundance of reads mapped to each genome was generated per sample. The statistical significance was measured using the Wilcoxon test.

### crAssphages *α*- and *β*-Diversities

From each subsampling exercise of 700,000 paired reads, the a-diversity metrics were determined, and the average of the 1,000 exercises was used as the final value for each sample. To contrast between groups, a Mann-Whitney-Wilcoxon test was performed. β-Diversity analyses were performed using the normalized reads-count table, using the Jaccard and Bray-Curtis metrics implemented in QIIME (v1.9).

### Association of crAssphage abundance with metadata variables and microbiota

We used the previously published biochemical and anthropometric data (Bikel et al., 2021) for comparison with the crAssphage abundances. The Associations between the crAssphage abundances and biochemical and anthropometric and microbiota were performed using the Spearman coefficient test in R using cutoffs of R > 0.7 and p-value < 0.005. For the abundance of microbiota, we used the data of the relative frequency of the 27 significant taxa previously reported between NW, O, and OMS in the same cohort (Gallardo-Becerra et al., 2020).

### Correlation of crAssphage abundance with their bacterial host

From the relative frequency of the microbiota analysis published for the same cohort (Bikel et al., 2021), we selected the abundance of the following taxa: Bacteroidetes, Bacteroidia, Bacteroidales, Prevotellaceae, *Prevotella, Collinsella, Bacteroides, Bacilli*, and *Eubacterium*, that were suggested as bacterial host of crAssphages. After that, we correlated them with the average abundance of crAssphage in all our samples using the Spearman coefficient test in R with cutoffs of R > 0.7 and p-value < 0.005.

## Acknowledgements

We thank Juan Manuel Hurtado Ramírez for informatics technical support and the computational server maintenance, as well as Filiberto Sanchez for experimental technical support. M.C.E., and L.G.B., thanks to the Doctoral Biochemical Sciences Program at IBT UNAM and CONACyT by doctoral fellowships C.V.U.: 887285 and 778192, respectively. This work was supported by the CONACyT grant Ciencia de Frontera 2019 263986 and by the DGAPA PAPIIT UNAM (IN215520).

